# Dorsomedial Striatum CB1R signaling is required for Pavlovian outcome devaluation in male Long Evans rats and reduces inhibitory synaptic transmission in both sexes

**DOI:** 10.1101/2024.05.01.592059

**Authors:** Catherine A. Stapf, Sara E. Keefer, Jessica M McInerney, Joseph F. Cheer, Donna J. Calu

## Abstract

Cannabinoid-1 receptor (CB1R) signaling in the dorsal striatum regulates the shift from flexible to habitual behavior in instrumental outcome devaluation. Based on prior work establishing individual, sex, and experience-dependent differences in Pavlovian behaviors, we predicted a role for dorsomedial striatum CB1R signaling in driving rigid responding in Pavlovian autoshaping and outcome devaluation. We trained male and female Long Evans rats in Pavlovian Lever Autoshaping (PLA). We gave intra-dorsomedial striatum (DMS) infusions of the CB1R inverse agonist, rimonabant, before satiety-induced outcome devaluation test sessions, where we sated rats on training pellets or home cage chow and tested them in brief nonreinforced Pavlovian Lever Autoshaping sessions. Overall, inhibition of DMS CB1R signaling prevented Pavlovian outcome devaluation but did not affect behavior in reinforced PLA sessions. Males were sensitive to devaluation while females were not and DMS CB1R inhibition impaired devaluation sensitivity in males. We then investigated how DMS CB1R signaling impacts local inhibitory synaptic transmission in male and female Long Evans rats. We recorded spontaneous inhibitory postsynaptic currents (sIPSC) from DMS neurons at baseline and before and after application of a CB1R agonist, WIN 55,212-2. We found that male rats showed decreased sIPSC frequency compared to females, and that CB1R activation reduced DMS inhibitory transmission independent of sex. Altogether our results demonstrate that DMS CB1Rs regulate Pavlovian devaluation sensitivity and inhibitory synaptic transmission and suggest that basal sex differences in inhibitory synaptic transmission may underly sex differences in DMS function and behavioral flexibility.

## INTRODUCTION

Impairments in behavioral flexibility occur across a range of mental health disorders including Substance Use Disorder, schizophrenia, Obsessive-Compulsive Disorder, and depression (Geramita et al., 2020; Jordan and Andersen, 2017; Kalivas and Volkow, 2005; Listunova et al., 2018; Simmler and Ozawa, 2019; Thoma et al., 2007). Preclinical studies suggest that sex and individual differences influence behavioral control when environmental conditions change from what is expected (Amaya et al., 2020; Bien and Smith, 2023; Keefer et al., 2020; Morrison et al., 2015; Nasser et al., 2015). Understanding the neurobiological underpinnings of individual and sex differences in behavioral flexibility may help to identify novel therapeutic targets for disorders of behavioral control.

Instrumental conditioning procedures in rats identified dorsal striatal regulation of behavioral flexibility, which involves dorsomedial and dorsolateral striatal (DMS, DLS) subdivisions. The shift from goal-directed to habitual behavior that occurs after extended instrumental experience is mediated by a shift from DMS to DLS control, respectively (Amaya and Smith, 2018; Dickinson et al., 1995; Gremel and Costa, 2013; Peak et al., 2019; Yin et al., 2005, 2004). Within the dorsal striatum (DS), multiple cell-types mediate the activity and output of each DS subregion. The majority of neurons in the DS are GABAergic medium spiny neurons (MSNs), the main type of projection neuron arising from the DS (Graveland and Difiglia, 1985). One of the most abundant receptor types in the DS is the Cannabinoid Receptor-1 (CB1R), which is a G-protein coupled receptor that is expressed pre-synaptically on glutamatergic terminals into the DS and locally on terminals of fast-spiking interneurons and MSNs (Gerdeman and Lovinger, 2001; Gerdeman et al., 2002; Lovinger and Mathur, 2012; Brian N Mathur et al., 2013; Wu et al., 2015). An instrumental outcome devaluation study shows that CB1R deletion in the orbitofrontal cortex-DS projection promotes devaluation sensitivity even during schedules of reinforcement that ordinarily drive habitual responding (Gremel et al., 2016), suggesting that that CB1R-mediated inhibition of synaptic inputs to DMS shift behavior towards rigid, devaluation insensitive instrumental actions.

We hypothesize that DMS CB1R signaling also biases behavior towards rigid devaluation insensitive Pavlovian behaviors. The sign-tracking model has uncovered considerable individual, sex and, experience-dependent differences in Pavlovian devaluation sensitivity (Flagel et al., 2009; Keefer et al., 2020; Kochli et al., 2020; Madayag et al., 2017; Pitchers et al., 2015), which suggest differences in endocannabinoid regulation of behavioral flexibility in the DMS. After limited training (<10 sessions) in Pavlovian lever autoshaping (PLA), in which an insertable lever cue predicts a food outcome, goal-tracking (GT) rats show sensitivity to outcome devaluation while sign-tracking (ST) rats do not (Keefer et al., 2020; Morrison et al., 2015; Nasser et al., 2015; Patitucci et al., 2016). After extended training (>10 session), both GT and ST rats show sensitivity to satiety-induced outcome devaluation (Keefer et al., 2020), an effect established in male rats. Female rats show increased levels of sign-tracking, or lever-directed approach during PLA compared to males (Hammerslag and Gulley, 2014; Keefer et al., 2022; King et al., 2020; Kochli et al., 2020; Madayag et al., 2017; Pitchers et al., 2015), suggesting they may be less sensitive to outcome devaluation even after extended training. In the present study, we use intracranial CB1R inverse agonist, rimonabant, to determine the role of DMS CB1R in mediating Pavlovian devaluation sensitivity in male and female rats.

Opposite to our prediction, we find that DMS CB1R signaling is necessary for flexible behavior in Pavlovian outcome devaluation. Based on this finding, and prior studies establishing that inhibition of DMS promotes rigid responding (Ragozzino et al., 2002; Yin et al., 2005), we hypothesized that CB1Rs located on GABAergic synapses onto MSNs in the DMS act to reduce inhibitory synaptic transmission. To test this, we measured the effect of CB1R activation on spontaneous inhibitory post synaptic currents (sIPSCs) in the DMS. In the slice electrophysiology studies, we include both males and females to investigate potential sex differences in DMS physiology. We found that male rats showed decreased sIPSC frequency compared to females, and that CB1R activation reduced DMS inhibitory transmission independent of sex.

## METHODS

### Subjects

For behavioral experiments, we used 68 Long Evans rats (33 Male, 35 Female; run as 5 cohorts) in the age range of 7-9 weeks old at the start of training for this study. All rats were double-housed upon arrival and then single-housed 24-48 hours after arrival. We performed all behavioral procedures during the dark phase of the light cycle. All rats had *ad libitum* access to standard laboratory chow and water before we food deprived them to maintain 90% of their baseline weight. We surgerized one cohort prior to any behavioral training and testing and surgerized the remaining cohorts after three days of training. There were no pre- or post-surgery differences in behavior between groups.

For slice electrophysiology experiments, we used 24 Long Evans rats (13 Male, 11 Female) in the age range of 9-15 weeks old at the time of slice electrophysiology recording. All rats were double housed upon arrival. These rats had ad libitum access to standard laboratory chow and water before we food deprived them 24 hours prior to slice electrophysiology recording.

We maintained all rats on a reverse 12hr:12hr light-dark cycle (lights off at 1000). We performed all procedures in accordance with the “Guide for the Care and Use of Laboratory Animals” (8^th^ edition, 2011, US National Research Council) and with approval by [Author University] Institutional Animal Care and use Committee (IACUC).

### Apparatus

We conduct behavioral experiments in identical operant chambers (25 × 27 × 30 cm; Med Associates) located in a separate room from the animal colony. An individual sound-attenuating cubicle with a ventilation fan surrounds each chamber. One wall contains a red house light and the opposing wall contains a food cup with photobeam detectors that rests 2 cm above the grid floor. A programmed pellet dispenser attached to the foodcup and dispensed 45 mg food pellets (catalog #1811155; Test Diet Purified rodent Tablet [5TUL]; protein 20.6%, fat 12.7%, carbohydrate 66.7%). We installed one retractable lever at 6cm above the grid floor on either side of the foodcup and we counterbalanced the lever side between subjects.

### Surgical Procedures

After three days of Pavlovian Lever Autoshaping training, we gave ad libitum access to food before we performed intracranial cannula placement surgery. We anesthetized 8-week-old rats with isoflurane (VetOne, Boise, ID, USA; 5% induction, 2-3% maintenance) then administered the pre-operative analgesic carprofen (5mg/kg, s.c.) and lidocaine (10mg/mL subdermal at incision site). We placed them in a stereotaxic frame (model 900, David Kopf Instruments, Tujunga, CA, USA) over a heating pad to maintain stable body temperature throughout surgery.

We implanted guide cannula (23G; PlasticsOne INC, Roanoke, VA, USA) bilaterally at an 8 degree angle and 1mm above the injection site into the DMS (coordinates from bregma -0.24 mm AP, ± 2.6 mm ML and -4.5 mm DV). We determined distance from bregma using the Paxinos and Watson rat brain atlas (Paxinos and Watson, 2006). Cannula were secured to the skull with jeweler’s screws and dental cement. At the end of surgery, we inserted dummy cannula into the guide cannula, which we only removed during infusion habituation and infusion test procedures. We moved rats to a recovery cage over a heating pad, administered carprofen analgesic at 24 hr, 48 hr and 72 hr post-surgery. We gave animals 1 week of recovery before resuming behavioral procedures.

### Pavlovian Lever Autoshaping Training

Prior to training, we exposed all rats to the food pellets in their home cage to reduce novelty to the food. Then we trained them in daily Pavlovian lever autoshaping sessions which lasted ∼ 26 minutes and included 25 trials of non-contingent lever presentations (conditioned stimulus; CS) and occurred on a VI 60 s schedule (50-70s). At the start of the session, the houselight turned on and remained on for the duration of the session. Each trial consisted of a 10 s lever presentation and retraction of the lever followed immediately by delivery of two 45 mg food pellets into the foodcup. At the end of the session, we returned rats to their cage and colony room. We trained rats in PLA first for 5 days to determine their tracking group, then continued training through 12 days following PLA testing.

### Pavlovian Lever Autoshaping Testing

We tested the effects of blocking DMS CB1R during reinforced Pavlovian Lever Autoshaping. We infused rimonabant (SR141716 1 µg/µL or 2 µg/µL; dissolved in 1:1:18 ethanol: emulphor: saline solution) or vehicle bilaterally into DMS at a rate of 0.5 µL/min over the span of 1 minute. We left the infusion cannula in place for an additional minute before slowly removing them and replacing the dummy cannula. We waited 10 min after infusion before placing rats into the behavioral chamber and running the lever autoshaping test. We infused all rats with vehicle, low (1 µg/µL) or high (2 µg/µL) dose of rimonabant across three days and we counterbalanced the dose across days.

### Satiety-Induced Outcome Devaluation Testing

After the 12^th^ training session, we gave rats two sessions of satiety-induced outcome devaluation. Rats had one hour of *ad libitum* access to 30 g of either their homecage chow (valued condition) or food pellets used during PLA training (devalued condition) in a ceramic ramekin. Within 15 min of the end of the satiation hour, we performed intra-DMS rimonabant infusions (2 µg/µL) as described in the previous section. We waited 10 min after the infusion before placing rats into the behavioral chamber and running the lever autoshaping test. Tests consisted of 10 non-rewarded lever presentations on VI 60s schedule (50-70s). Immediately after each test, we gave rats a 30 min food choice test in their homecage which included 10 g of homecage chow and 10 g of food pellets in separate ceramic ramekins to confirm satiety was specific to the outcome they had been fed before the test session. We retrained rats on 25 reinforced trials on a separate day between devaluation probe tests.

### Brain Slice Preparation for Slice Electrophysiology

We anesthetized rats with isoflurane then perfused with chilled N-Methyl-D-Glucamine (NMDG)-modified artificial cerebrospinal fluid (NMDG-aCSF; in mM; 92 NMDG, 20 HEPES, 25 Glucose, 30 NaHCO3, 1.3 NaH2PO4, 2.5 KCl, 5 Sodium Abscorbate, 3 Sodium Pyruvate, 2 Thiourea, 10 MgSO4, 0.5 CaCl2) that had been bubbled with carbogen (95% oxygen, 5% carbon dioxide). We collected coronal sections from the DMS (350µM) while the brain was mounted on the cutting stage and submerged in chilled, carbogen-bubbled NMDG-aCSF, using a Leica VT 1200 vibratome. We incubated slices in carbogen-bubbled, 40° NMDG solution for 5-8 minutes then transferred slices to room temperature, carbogen-bubbled HEPES holding solution (in mM; 92 NaCl, 20 HEPES, 25 Glucose, 30 NaHCO3, 1.3 NaH2PO4, 2.5 KCl, 5 Sodium Abscorbate, 3 Sodium Pyruvate, 2 Thiourea, 1 MgSO4, 2 CaCl2). We waited 1 hour before making the first recordings. Sections remained in the holding solution until electrophysiological recordings were performed.

### Recordings and Bath Application of Drug

We visualized cells in the DMS using an Olympus BX50 light microscope. We recorded spontaneous IPSCs (sIPSC) using borosilicate, fire-polished glass pipettes with resistance in the 3-5 MΩ range. We pulled pipettes with a Narshige PC-100 pipette puller and filled them with a CsCl-based internal solution (in mM; 150 CsCl, 10 HEPES, 2 MgCl2*H2O, 0.3 Na-GTP, 3 Mg-ATP, 0.2 BAPTA). We recorded from hemisected slices that were constantly perfused with 37° carbogen-bubbled artificial cerebrospinal fluid (aCSF; in mM; 126 NaCl, 25 NaHCO3, 11 Glucose, 1.2 MgCl2*H2O, 1.4 NaH2PO4, 2.5 KCl, 2.4 CaCl2), containing blockers of AMPA (DNQX, 20µM) and NMDA (AP5, 50µM). We perfused the recording chamber with a basic Longer Pump BT100-2J peristaltic pump. We also recorded from slices submerged in a bath containing DMSO (0.065%) and 2-hydroxypropyl-beta-cyclodextrin (0.006%). We clamped cells at -60 mV using a Molecular Devices Multiclamp 700B amplifier and digitized recordings with a Molecular Devices Axon Digidata 1550B digitizer. We used Molecular Devices Clampex 10.7 software for data acquisition. We excluded recordings when sIPSC baseline was below -200 pA, series resistance was >40 MΩ, or series resistance changed >20% throughout the course of the experiment.

### Measurements

For training and devaluation probe tests, we recorded the number and duration of foodcup and lever contacts, the latency to contact, and the probability during the 10 s CS (lever) period. On trials with no contacts, a latency of 10s was recorded. To determine tracking group, we used a Pavlovian Conditioned Approach (PavCA) analysis (Meyer et al., 2012) which quantifies behavior along a continuum where +1.00 indicates behavior is primarily lever directed (sign-tracking) and -1.00 indicates behavior is primarily foodcup directed (goal-tracking). PavCA scores are the average of three separate scores: the preference score (lever contacts minus foodcup contacts divided by the sum of these measures), the latency score (time to contact foodcup minus the time to contact lever divided by 10 s (duration of the cue)) and the probability score (probability to make a lever contact minus the probability to make a foodcup contact across the session). We use the PavCA score from the 5^th^ day of training to determine an individual’s tracking group as follows: sign-trackers (ST) have a PavCA sore +0.33 to +1.00, goal-trackers (GT) have a PavCA score -1.00 to -0.33, intermediates (INT) have scores ranging from -0.32 to +0.32. Rats in goal- and intermediate tracking groups were combined into a single GT/INT group as they show similar patterns of outcome devaluation in other studies (Keefer et al., 2020). On day 6, we were unable to record latency data for 6 rats and only retained lever and foodcup contacts for these rats. Preference score was used in place of PavCA for rats on this day.

For devaluation probe tests, we also report total approach (the sum of food cup and lever contacts during the 10 s CS period) and individual contact measurements. We recorded consumption on the test days and calculated the amount of pellet or chow consumed in grams during the satiety hour and during the 30 min choice test.

We processed sIPSC traces using the template search function in Molecular Devices Clampfit 10.7 software to determine event peak amplitude and event peak start time. We report these measurements in each experiment: Amplitude, calculated as the peak amplitude of an event and averaged across each recording; Frequency, calculated as the number of events per recording divided by the duration of the recording in seconds; Interevent Interval, calculated as the inverse of the time (in seconds) between the peak of an event and the peak of the event prior and represented through a cumulative frequency distribution.

### Histology

At the end of behavioral experiments, we deeply anesthetized rats with isoflurane and transcardially perfused 100ml of 0.1M sodium phosphate buffer (PBS), followed by 200ml of 4% paraformaldehyde (PFA) in PBS. We removed brains and post-fixed them in 4% PFA over night before we transferred them to 30% sucrose in dH2O for 48-72 hr at 4 °C. We rapidly froze brains in dry ice before storing them in -20 °C until slicing. We sliced brains with the Leica Microsystems 1850 cryostat to collect 40 µm coronal sections in three series through the cannula placements in the DMS. We mounted sections onto gel-coated slides and then stained with cresyl violet before coverslipping with Permount. We examined placements under a light microscope for confirmation of cannula placement in the DMS (Fig. 2B). We excluded 11 rats due to cannula placements being outside the region of interest.

### Experimental Design and Statistical Analysis

We analyzed behavioral data using SPSS 29.0 statistical software (IBM) or Prism (Graphpad software) with mixed-design repeated measures analysis of variance (ANOVA) or paired t tests, where applicable. Significant main and interaction effects (p<0.05) were followed by post-hoc repeated-measures ANOVA or Bonferroni comparisons. Analyses included between subject factors of Tracking (ST, GT/INT) Sex (male, female) and Treatment (vehicle, rimonabant) and within-subject factors of Session (1-12), Outcome Value (Valued, Devalued), or Outcome (Nonsated, Sated).

For slice electrophysiology experiments, data are represented as mean ± standard error or presented as cumulative frequency distribution plots. We performed independent samples student’s t-test, two sample Kolmogorov-Smirnov tests, or Kruskal-Wallis tests with Dunn’s post hoc comparisons as appropriate using either SPSS or Prism. We analyzed mean amplitude and mean frequency data using independent samples t-tests between males and females. We analyzed the cumulative frequency distribution of interevent interval between males and females using a Kolmogorov-Smirnov test and reported the effect size using Hedges’ *g.* We analyzed the cumulative frequency of interevent intervals between DMSO and WIN conditions in the bath and between males and females using the Kruskal-Wallis test with Dunn’s post hoc comparisons. Analysis included within-subject variable of Bath (pre-WIN, post-WIN) and between-subject variable of Sex (Male, Female). We removed two data points, one from each Sex, based on results from Grubb’s Test for Outliers.

## RESULTS

### Acquisition of Pavlovian Lever Autoshaping differs due to Tracking and Sex

We trained rats for 12 days in Pavlovian Lever Autoshaping in which an insertable lever cue predicts food pellet delivery. We used the Pavlovian Conditioned Approach Index (PavCA) on the 5^th^ session of training to determine tracking groups (Fig. 1A). We then used a mixed design repeated measures ANOVA with between subject factor of Tracking (ST, GT/INT) and within subject factor of Session (1-12). Consistent with group assignments, ST rats show more lever directed behavior than GT/INT rats (main effect Tracking; F(1,53) = 49.293, p=<0.001). Consistent with prior studies (Villaruel and Chaudhri, 2016; Bacharach et al., 2018; Keefer et al., 2020) showing that GT and intermediate (GT/INT) rats shift away from food-cup approach and towards lever approach with extended training, we observe a main effect of Session for PavCA Index, F(11,583)=106.292, p<0.001, and a Session x Tracking (ST, GT/INT) interaction, F(11,583)=13.909, p<0.001). Next, we examined whether there were sex differences in the acquisition and expression of Pavlovian approach behaviors (Fig. 1B). We used a similar statistical approach with between-subject factor of Sex (Male, Female) and within subject factor of Session (1-12) and found a Session x Sex interaction for PavCA Index, F(11,605)=1.823, p=0.047). We analyzed PavCA indices between males and females using independent samples t-tests for each day. While males and females show similar PavCA indices during initial acquisition, female rats showed more sign-tracking, via a higher PavCA Index, than males with extended training (day 8, t(55)=-1.754, p=0.043; day 9,t(55)=-2.007, p=0.025). However, there were no sex differences in responding on the last day of training (PavCA Index; t(55)=-1.099, p=0.277), prior to testing in outcome devaluation.

**Figure 1.**
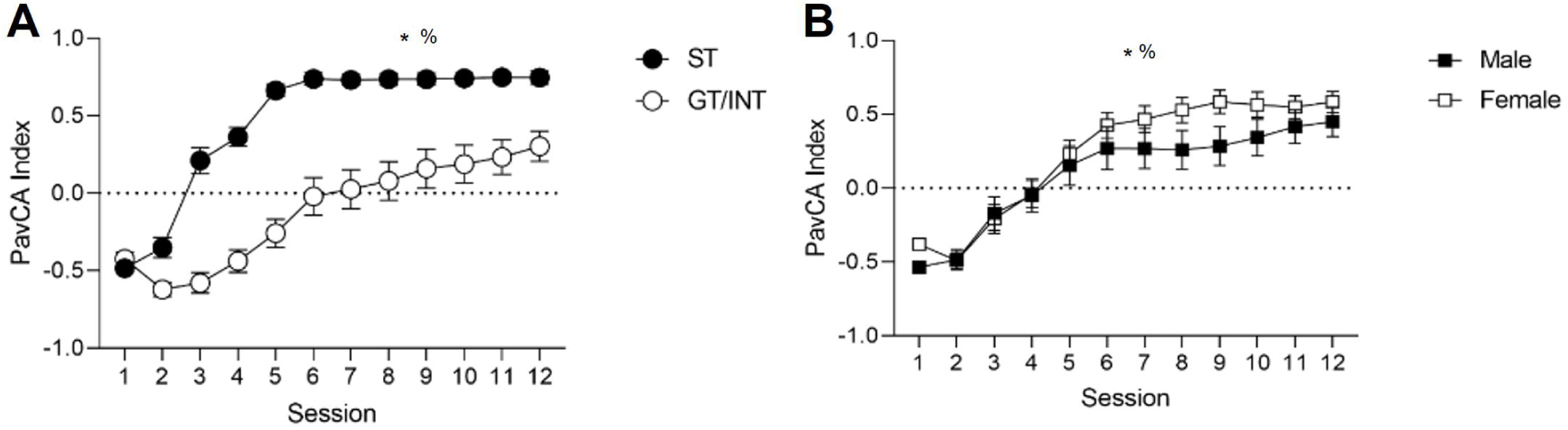
Acquisition of a Pavlovian Conditioned Approach differs by Tracking and Sex. *A,* PavCA Index mean ± SEM for ST and GT/INT (collapsed on sex) that acquire individual differences in conditioned responding in PLA task. *Main effect of Session. %Significant Session X Tracking interaction. *B,* PavCA Index mean ± SEM for Male and Female rats (collapsed on tracking) that acquire conditioned responding in a PLA task.

### Intra-DMS inhibition of CB1R signaling prevents outcome devaluation but does not affect Pavlovian Approach during non-sated, reinforced sessions

We tested rats using a within-subject satiety-induced outcome devaluation procedure in which they were sated on the training pellet (devalued) or homecage chow (valued) just prior to brief PLA test sessions under extinction conditions. Prior to test sessions we gave intra-DMS vehicle or CB1R inverse agonist, rimonabant, injections to determine the effects of inhibiting DMS CB1R signaling on devaluation sensitivity of Pavlovian approach. To examine how this manipulation generally affects Pavlovian devaluation sensitivity across all rats, we analyzed total approach which is the sum of lever and foodcup contacts during the 10 s cue presentation. We compared responding during the valued (chow sated) versus devalued (pellet sated) conditions using a mixed design repeated measures ANOVA with between subject factor of Treatment (Vehicle, Rimonabant) and within subject factor of Outcome Value (Valued, Devalued). Figure 2A shows the performance of all rats that received either intra-DMS vehicle or rimonabant infusions during the outcome devaluation probe test. We found a main effect of Outcome Value (F(1,49)=5.558, p=0.022) and an Outcome Value x Treatment interaction (F(1,49)=6.663, p=0.013), indicating that intra-DMS rimonabant impaired Pavlovian devaluation sensitivity across all rats. Under vehicle conditions, rats decreased total approach when sated on the training pellet (devalued state) compared to when they were sated on the homecage chow (valued state). In contrast, with intra-DMS rimonabant infusions, rats showed a similar amount of Pavlovian approach in the valued and devalued states. These results suggest a divergent endocannabinoid mechanism for mediating Pavlovian outcome devaluation in which DMS CB1R promote flexibility, in contrast to prior studies suggesting that CB1R signaling promotes rigid responding in instrumental settings (Navarro et al., 2001; Hilário et al., 2007; Gremel et al., 2016). Figure 2B shows the approximate location of intra-DMS infusions.

**Figure 2.**
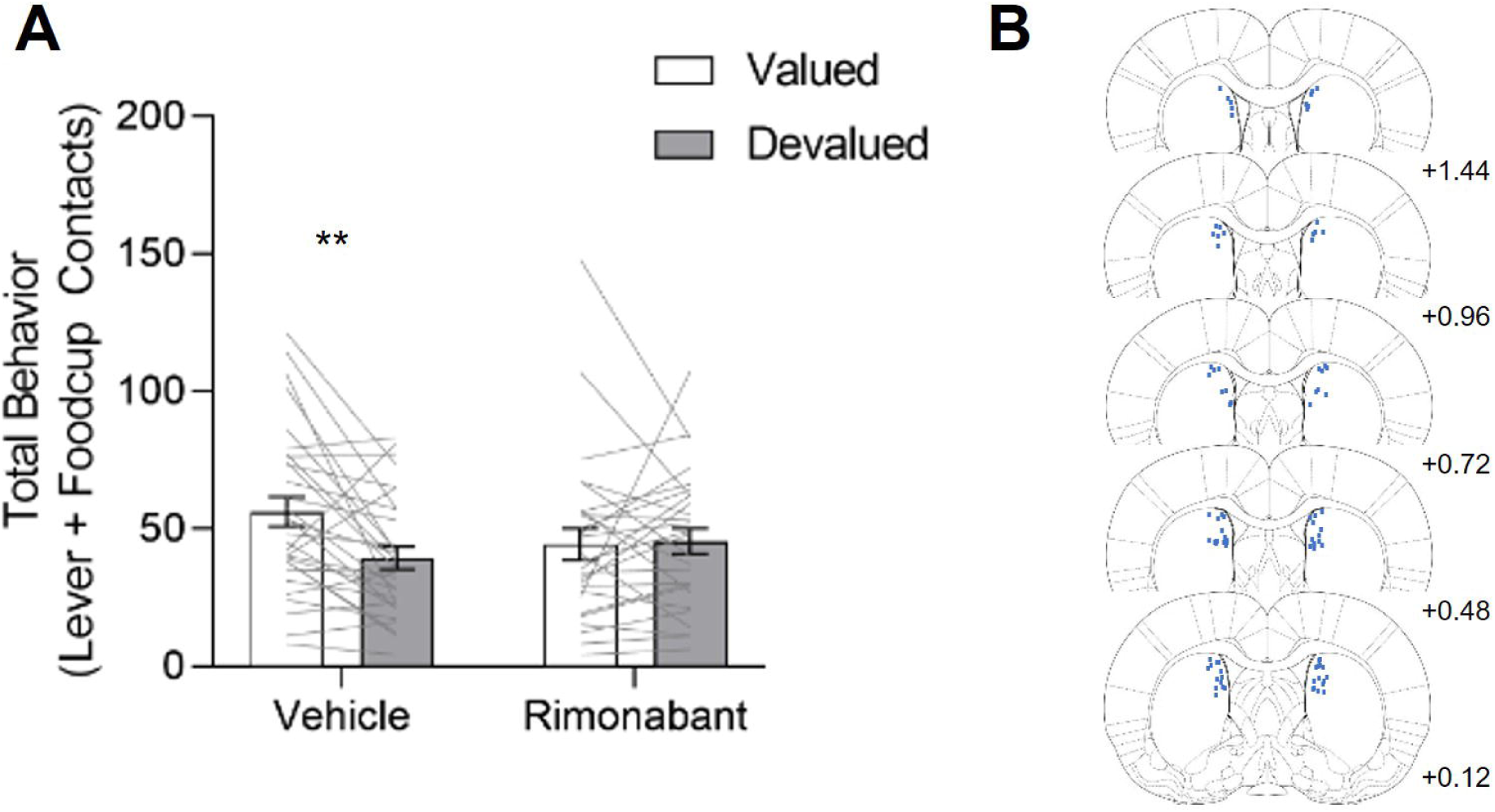
Intra-DMS Rimonabant prevents sensitivity to Pavlovian Outcome Devaluation. Data are represented as within-subject individual data (lines) and group data (bars; mean ± SEM). Rats received intra-DMS injections of either vehicle (left) or rimonabant (right) 10 minutes prior to probe test. ***A,*** Total behavior (sum of lever and food cup contacts) in outcome devaluation across all rats. We observed a main effect of Outcome Value and a significant Outcome Value X Treatment interaction. ***B,*** Coronal sections (in mm) depicting the location of DMS injector tips for rimonabant infusion. **p<0.025

Considering the established individual differences in devaluation sensitivity in Pavlovian autoshaping (Keefer et al., 2020; Morrison et al., 2015; Nasser et al., 2015; Patitucci et al., 2016; Smedley and Smith, 2018), we added Tracking and Sex as between-subject factors in this analysis. We observed an Outcome Value x Treatment x Sex x Tracking interaction (F(1,49)=4.545, p=0.038)) which points to differences in the effects of treatment on devaluation sensitivity that differ by Sex and/or Tracking. In male rats we observed a main effect of Outcome Value and an Outcome Value x Treatment interaction (Fig. 3A, Value: F(1,25)=6.084, p=0.021; Value X Treatment: F(1,25)=6.440, p=0.018). Bonferroni post hoc comparisons confirm that under vehicle conditions, male rats were sensitive to outcome devaluation (t(53)=3.905, p<0.007) responding more to cues in valued than in devalued conditions. We observed that intra-DMS rimonabant impaired devaluation sensitivity in male rats, as they responded similarly between valued and devalued conditions (t(53)=0.0534, p>0.999). In female rats, we did not observe any significant main effects or interactions (Fig. 3B; *Fs*<0.890, *ps*>0.353), indicating they were not sensitive to Pavlovian outcome devaluation; thus, we could not evaluate treatment effects on this behavior.

**Figure 3.**
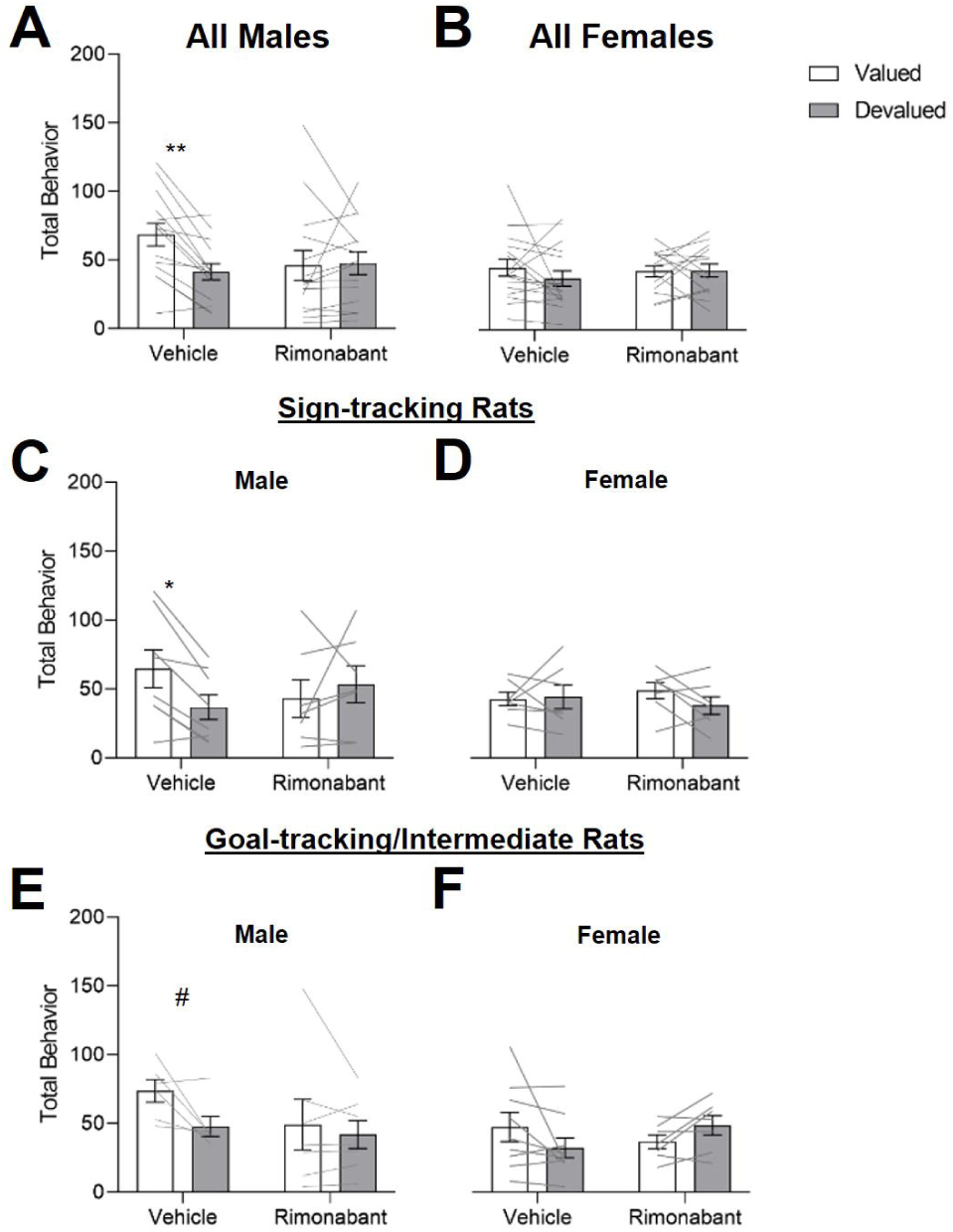
Male, but not female, rats are sensitive to Pavlovian Outcome Devaluation, and this sensitivity is blocked by intra-DMS Rimonabant regardless of Tracking type. Data are represented as within-subject individual data (lines) and group data (bars; mean ± SEM). Rats received intra-DMS injections of either vehicle (left) or rimonabant (right) 10 minutes prior to probe test. ***A,*** In Male rats, we observed a significant main effect of Outcome Value and a significant Outcome Value X Treatment interaction. ***B***, In Female rats, we did not observe any significant main effects or interactions. ***C, D***, In ST rats, we observe a significant Outcome Value X Treatment X Sex interaction on total behavior. ***C,*** In Male ST rats we observed a significant Outcome Value X Treatment interaction. ***D,*** In ST Female rats there were no main effects or interactions. We then performed a parallel analysis in our GT/INT rats. ***E, F*** ln GT/INT rats, we observe an Outcome Value X Treatment interaction, but no interaction with Sex. #p=0.067 *p<0.05 **p<0.025

In a prior study using male rats, it was established that initially devaluation insensitive ST rats become devaluation sensitive after extended training (Keefer, 2020). The present study replicates this finding and shows that under vehicle conditions, male ST rats are sensitive to outcome devaluation (Fig. 3C, Bonferroni post-hoc; t(13)=2.679, p=0.037). Here we use both sexes and identify an Outcome Value x Treatment x Sex interaction in ST rats (F(1,24)=6.210, p=0.020), suggesting potential sex differences in devaluation sensitivity and/or effects of CB1R signaling inhibition. First, we confirmed the Outcome Value x Treatment interaction that was observed overall (Fig. 2A) is also observed in male ST rats (Fig. 3C, F(1,12)=5.063, p=0.044). Post-hoc analyses confirmed that intra-DMS rimonabant injections impaired devaluation sensitivity in male rats with similar levels of Pavlovian approach for valued and devalued conditions (t(13)=0.9205, p=0.7482). We found similar trends for male ST rats in lever contacts (the dominant response of ST rats) during outcome devaluation (Fig. 3-1A), in which there was a significant Outcome Value X Treatment interaction (F(1,13)=4.810, p=0.047) but post hoc tests did not reach significance even for the vehicle condition (t<2.484, p>0.0548). As expected, we observed no significant effects when analyzing male ST foodcup contacts (Fig. 3-2A). In contrast to males, female ST rats showed similar levels of responding in all probe tests and intra-DMS rimonabant had no effects (Total Behavior, Fig. 3D, *Fs*<1.236, *ps*>0.288; Lever, Fig. 3-1B; Foodcup, Fig. 3-2B).

Consistent with prior studies, male GT/INT rats were sensitive to outcome devaluation after extended training (main effect of Outcome Value (Fig. 3E; F(1,11)=5.203, p=0.043). In contrast to the ST group, we observed no main effects or interactions with Sex in GT/INT group. Despite this, we performed parallel analyses and found a marginal devaluation effect under vehicle condition in male GT/INT rats (t(11)=2.425, p=0.0675). For GT/INT the dominant response is food cup contacts, and for this measure there was a significant Outcome Value X Treatment interaction (Fig. 3-2C; F(1,11)=7.279, p=0.0207) and post hoc analysis revealed that under vehicle conditions, male GT/INT rats were sensitive to outcome devaluation (t(11)=2.872, p=0.0304) which was not the case with intra-DMS rimonabant (t(11)=0.8692, p=0.8066). We observed no significant differences when analyzing lever contacts alone (Fig. 3-1C). Female GT/INT rats showed a significant Outcome Value X Treatment interaction for total behavior (Fig. 3F; F(1,14)=5.100, p=0.040) that was driven by opposite patterns of behavior for the two treatments, however differences between value conditions did not reach significance for either treatment (vehicle, valued vs. devalued, t(14)=1.907, p=0.1545, rimonabant, valued vs. devalued (t(14)=1.329, p=0.410). We found a similar Outcome Value X Treatment interaction when looking at female GT/INT lever contacts alone (Fig. 3-1D; F(1,14)=4.953, p=0.043) but no significant interactions for food cup contacts (Fig. 3-2D); however, none of the post hoc analysis for these measures reached significance in female GT/INT rats.

Altogether, these results point to sex differences in Pavlovian outcome devaluation sensitivity and to treatment effects on Pavlovian devaluation sensitivity in male rats. Male rats are sensitive to devaluation after extended training, while female rats are not. The effects of intra-DMS CB1R blockade on devaluation sensitivity in male rats are consistent across tracking groups but are response specific. In male ST rats this sensitivity is driven by lever contacts, while in male GT/INTs, this sensitivity is driven by food cup contacts.

These effects of DMS CB1R signaling inhibition were specific to the satiety-specific outcome devaluation test. We found no difference in responding between intra-DMS vehicle and rimonabant groups during a non-sated, non-reinforced Pavlovian lever autoshaping test of the same duration (10 trials, Fig. 2-1A; Sex x Treatment x Response (lever, foodcup), *Fs*<0.479, *ps*>0.493). This suggests that intra-DMS rimonabant treatment effects on Pavlovian approach emerge only after outcome-specific satiety. The observed effects are also not due to differences in consumption between male and female rats during the 1-hour satiation period. To account for body weight differences between male and female rats of the same strain and age, we normalized the amount (g) of food consumed (either for the satiation period or post-probe choice test) to each rat’s average body weight across both days of outcome devaluation tests (Council, 1995; Lenglos et al., 2013). We found no difference in the amount of food consumed during the satiation period prior to the probe test (g/bw chow Mean: Male, 0.032, SEM ±0.002; Female, 0.031, SEM ±0.002; g/bw pellet Mean: Male, 0.039, SEM ±0.002; Female, 0.036, SEM ±0.002; *Fs*<1.153, *ps*>0.288). To confirm devaluation of the sated food, we gave rats a post-probe choice test between the chow and training pellets (Fig. 2-1B) immediately after the end of the outcome devaluation probe test. Rats consumed less of the food when they were sated on and more of the alternative food, verified by a main effect of Outcome (F(1,45)=8.134, p<0.007) and this did not differ by sex or treatment (*Fs*<1.790, *ps*>0.187).

The observed effects of inhibiting CB1R signaling were also not evident during non-sated, reinforced P vlovian lever autoshaping sessions. We tested the effect of intra-DMS rimonabant infusion on a subset of rats (N=12) during non-sated, reinforced PLA sessions and found no significant difference between vehicle, low (1µg/µl) or high dose (2µg/µl) of rimonabant on lever presses (Fig. 2-2A; *Fs*<1.972, *ps*>0.198) or on foodcup contacts (Fig. 2-2B; *Fs*<1.078, *ps*>0.329) across sex or tracking. We did observe a significant main effect of Sex for lever contacts (F(1,10)=5.395, p=0.043), which is in line with acquisition data, during which females showed more sign-tracking. Overall, rimonabant inhibition of DMS CB1R signaling did not impact conditioned approach under reinforced conditions.

### Baseline spontaneous IPSC recordings in DMS neurons differ between male and female Long Evans rats

Based on the sex differences in behavioral flexibility, we predicted that there may be differences in inhibitory synaptic transmission in the DMS, in which male rats may show reduced inhibitory synaptic transmission. We recorded spontaneous IPSCs from cells in the DMS in males and females (Fig. 4A). We examined the mean amplitude (absolute value), the mean frequency, or total events across the duration of the recording, and the cumulative frequency distribution for interevent interval, or the time between event peaks, during 5-min recordings. We found that there is no difference in the mean amplitude of DMS sIPSCs between males and females when slices are perfused with an aCSF bath (Fig. 4B, t= -1.226, p=0.239). However, we did see a difference in both the frequency and interevent interval. Male rats show a lower frequency as compared to females (Fig. 4C, t= - 2.561, p=0.022) and a larger inter-event interval (Fig. 4D, Kolmogorov-Smirnov test, D=0.2498, p<0.0001, Hedge’s *g* = 0.426). This difference in frequency and inter-event interval of sIPSCs suggests that male rats show less inhibitory synaptic transmission onto recorded DMS neurons than females.

**Figure 4:**
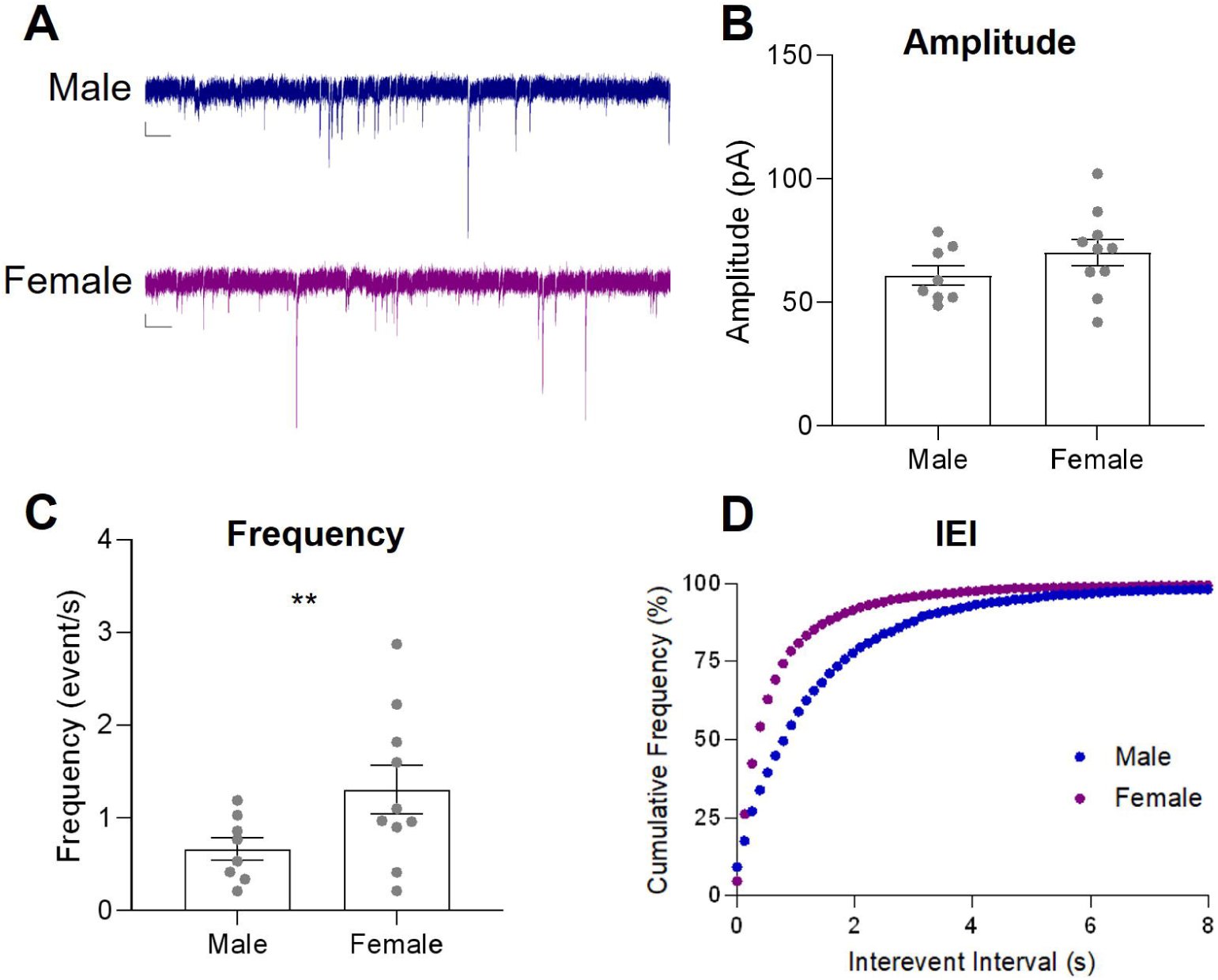
sIPSCs In DMS cells show reduced frequency and larger Inter-event Intervals In melee as compared to females. ***A,*** Representative sIPSC recordings from DMS cells in aCSF bath of male (blue, n=8) and female (purple, n=9) Long Evans Rats. Scale bare: 20 pA and 1 sec. Data are presented as mean ± SEM. ***B,*** Mean Amplitude ***C,*** Mean Frequency ***D,*** Cumulative Frequency Plot of Interevent Interval. **p<0.025

### WIN 55, 21-2 bath application changes sIPSC measures in both males and females relative to DMSO bath application

We hypothesized that activation of DMS CB1R would reduce inhibitory synaptic transmission in male rats and included females to investigate if there are sex differences in the effect of CB1R manipulation on sIPSCs in the DMS. We recorded sIPSCs from DMS cells for 5 mins at baseline and following a 10-minute bath application of a CB1R agonist, WIN 55,212-2 (WIN; 10µM, Fig. 5A). We found that there were no differences in the mean amplitude of sIPSCs due to WIN or Sex (Fig. 5B, *Fs*<1.182, *ps*>0.290). However, we did see differences in frequency and inter-event interval (Fig. 5C,D). We found a main effect of WIN for frequency (F(1,19)=6.306, p=0.021) but no main effect or interaction with Sex (*Fs*<0.825, p>0.375). We found that application of WIN shifted the interevent interval cumulative distribution curves to the right (Kruskal Wallis, H=1359, p<0.001) and post hoc comparisons confirmed that this occurred for both male and female rats (DMSO vs. WIN; Dunn’s comparisons; male, p<0.0001, Hedges’ *g* = 0.2085; female, p<0.0001, Hedges’ *g =* 0.2291). This rightward shift suggests that WIN increases the interevent interval in both sexes. Application of WIN in the bath caused a reduction in the frequency of inhibitory events and an increase in the inter-event interval across all rats, suggesting that CB1R located on presynaptic inhibitory inputs suppresses release of GABA in the DMS.

**Figure 5:**
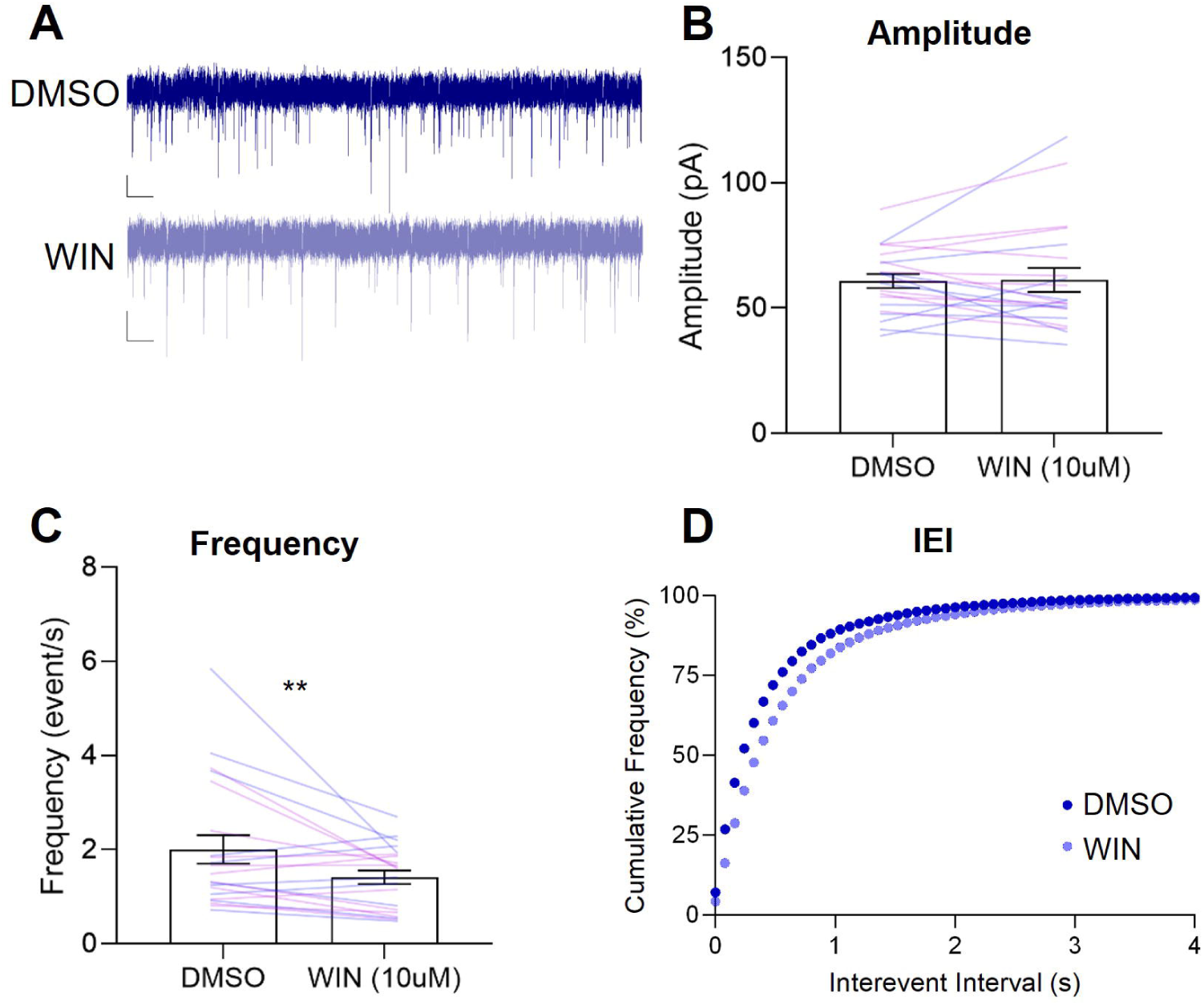
Activation of CB1R by WIN reduces sIPSC frequency and increases sIPSC interevent interval, regardless of Sex. ***A,*** Representative sIPSC recordings from DMS cells pre- (blue) and post-WIN (light blue) bath application for Long Evans Rats (n=21). Scale bars: 20 pA and 1 sec. Data are presented as mean ± SEM. ***B,*** Mean Amplitude with individual data for males (blue) and females (purple). ***C,*** Mean Frequency with individual data for males and females ***D,*** Cumulative Frequency Plot of Interevent Interval of sIPSCs. **p<0.025

## DISCUSSION

In the current studies we investigated the role of DMS CB1R signaling in Pavlovian outcome devaluation and regulation of inhibitory synaptic transmission. We found that after extended training in PLA, males are sensitive to outcome devaluation, while females are not and that DMS CB1Rs were necessary for the devaluation sensitivity in males. Slice electrophysiology studies revealed that male rats showed a reduced frequency of inhibitory events in the DMS as compared to females but that activating DMS CB1Rs reduced the prob bility of GABA release similarly in both sexes.

The current results align with prior research that established significant individual-, experience-, and sex-dependent differences in Pavlovian devaluation. Consistent with previous studies (Pitchers et al., 2015; Keefer et al., 2022; Kochli et al., 2020), we found that female rats showed more lever-directed behaviors than males during extended training, but this difference diminished before testing in outcome devaluation (Fig. 1B). Under vehicle conditions, we replicated prior findings that male rats show devaluation sensitivity after extended training in PLA (Keefer, 2020). We extended this work to include females, for which we do not observe devaluation sensitivity after extended training (Fig. 2). These results echo the findings of other studies that indicate females are less sensitive to instrumental and Pavlovian devaluation (Bien and Smith, 2023; Quinn et al., 2007; Schoenberg et al., 2019; Sood and Richard, 2023). Additionally, further analyses into either lever or foodcup contacts revealed that male rats were sensitive to devaluation for their preferred response of their tracking group. ST male rats reduced lever contacts when the outcome is devalued (Fig. 3-1A), while GT/INT male rats reduce foodcup contacts (Fig. 3-2C) as has been shown previously in studies that examine Pavlovian outcome devaluation after extended training (Keefer, 2020; Keefer et al., 2022).

At first, we predicted that dorsal striatal CB1R signaling would promote rigid, or habitual, behaviors as has been shown for instrumental outcome devaluation (Gremel et al., 2016). However, our study suggests that CB1Rs in DMS promote behavioral flexibility in male rats, running counter to this established understanding. There are several factors that contribute to the divergence of results including species differences, the use of Pavlovian versus instrumental devaluation procedures and the subregion-specific effects of experimental manipulations. This prior study trained CB1R flox mice in both random-ratio (RR) and random interval (RI) schedules of instrumental reinforcement and generated an OFC-DS specific CB1R knockout. Competing action-outcome and stimulus-response associations mediate instrumental devaluation, and studies show that goal-directed behaviors shift to habit with extended training or with RR schedules of reinforcement (Adams, 1982; Adams and Dickinson, n.d.; Gremel et al., 2016). This is not the case with Pavlovian behaviors that are sensitive to devaluation even after extended training (Holland, 1998; Keefer et al., 2020), suggesting stimulus-outcome associations support adaptive reward seeking despite overtraining. Thus, differences in Pavlovian and instrumental processes may, in part, underline divergent findings between studies. Another possibility is methodological differences in the way CB1R were manipulated between studies. CB1R deletion in the OFC-DS projection promoted “goal-directed” devaluation sensitivity even during schedules of reinforcement that ordinarily drive “habitual” devaluation insensitivity (Gremel et al., 2016). Our current behavioral study inhibits CB1R signaling indiscriminately-likely affecting both inhibitory and excitatory synaptic transmission- rather than specifically on glutamatergic OFC afferents to the dorsal striatum, as in the prior study. Never-the-less, prior work has shown that systemic activation of CB1Rs promotes rigid responding (Hilário et al., 2007; Nazzaro et al., 2012) and while both DLS and DMS express CB1Rs (Hohmann and Herkenham, 2000; Fusco et al., 2004; Van Waes et al., 2012), more of the CB1R work within subregions of the DS has focused on the DLS. The DLS does express CB1R more densely than DMS, thus, it is possible that off-target effects impacted DLS function, an area with high CB1R density (Hohmann and Herkenham, 2000; Fusco et al., 2004; Van Waes et al., 2012) and this could confound our results. We think this is unlikely given the volume of rimonabant injected (0.5 µL per hemisphere) and our ex vivo confirmation of reduced inhibitory synaptic transmission with CB1R activation in the DMS. The current targeting of DMS, as compared to DLS, may in part explain why our results diverge from observations that dorsal striatal CB1Rs support rigid responding via inhibition of glutamatergic inputs and our findings fit within the context of the DMS’ role of biasing behavior towards “goal-directed” responding (Yin et al., 2005; Corbit and Janak, 2010; Gremel and Costa, 2013; Li et al., 2022).

These prior studies established that the DMS supports flexible, goal-directed instrumental conditioned responding. Reducing the activity of the DMS through lesion or pharmacological inhibition impairs flexible responding in a variety of tasks. To be interpreted in this conceptual framework, our behavioral pharmacology results suggest that CB1R signaling disinhibits the DMS, and thus, reducing CB1R signaling has a net inhibitory effect on DMS, resulting in impaired “goal-directed” Pavlovian devaluation sensitivity. Based on this interpretation, we hypothesized that DMS CB1R signaling at GABAergic inputs to DMS medium spiny neurons reduces inhibitory transmission in the area, allowing DMS activation to promote flexible responding in Pavlovian devaluation.

Our slice electrophysiology studies focused on inhibitory synaptic currents to investigate this hypothesis. At baseline, we found that males showed reduced inhibitory events as compared to females (Fig. 4). Within the above framework of striatal contributions to goal-directed and habitual control of behavior, lower levels of DMS inhibitory transmission (as seen in males) would promote flexibility and higher levels of inhibitory transmission (as seen in females) would prevent the expression of outcome devaluation, consistent with our devaluation findings in male and female rats, respectively. While we did not confirm the identity of the cells we recorded from, approximately 90% of cells across the dorsal striatum are medium spiny neurons (MSNs), the main type of projection neurons arising from the striatum (Graveland & Difiglia, 1985). Due to their abundance, we are likely to be recording from MSNs in the DMS. Multiple studies have shown that intact female rats and males treated with estradiol have increased striatal MSN excitability (Tansey et al., 2008; Dorris et al., 2015; Cao et al., 2018; Proaño et al., 2018) and estradiol decreases GABA release (Schultz et al., 2009). However, these studies are not specific to the DMS. Additionally, some studies have shown lower numbers of GABAergic neurons in males compared to females (Ovtscharoff et al., 1992), which may explain reduced inhibitory synaptic transmission in males. However, there are many types of GABAergic cells in the DMS. GABAergic Medium Spiny Neurons (MSNs) are the main projection neuron of the DMS, and they also project locally to other MSNs (Wilson and Groves, 1980; Somogyi et al., 1981; Graveland and Difiglia, 1985; Tunstall et al., 2002; Czubayko and Plenz, 2002; Burke et al., 2017). There are also multiple GABAergic interneuron types, predominately Parvalbumin positive fast-spiking interneurons (FSIs) and somatostatin interneurons (SOM). In fact, a study focusing on sex differences in the number of interneurons shows that some GABAergic interneurons are more dense in males than females (FSIs) while other interneurons are less dense in males than females (Van Zandt et al., 2024). Thus, further work must be done to isolate inhibitory synaptic transmission from these different sources and better understand sex differences in the DMS with cell-type specificity.

We show that CB1R activation reduces the frequency of inhibitory events regardless of sex. This should be interpreted with caution, as we only tested a single dose of the CB1R agonist. We applied WIN 55,212-2 at a concentration of 10 µM, which is a high concentration for bath application. Other studies use much lower doses and have seen sex differences in other brain regions (Tabatadze et al., 2015; Ferraro et al., 2020). Both males and females express CB1R in the dorsal striatum and males express CB1R more densely in the striatum and other brain regions than females (Laurikainen et al., 2019; Liu et al., 2020). Thus, it is possible that application of WIN at a lower dose may reveal more sensitivity to CB1R manipulation in males due to this higher concentration of receptors. Another caveat of these electrophysiological findings is that rats we recorded from did not have any behavioral training. It is possible that behavioral experience alters DMS inhibitory tone or changes DMS activity, as has been shown in other studies examining DMS activity after extended training or under different schedules of reinforcement (Fanelli et al., 2013; Gremel and Costa, 2013; Vandaele et al., 2019).

CB1Rs are located on multiple cell types in the dorsal striatum so further work must been done to identify the cell-type that supports Pavlovian flexibility in male rats. One notable possibility is the parvalbumin positive FSIs. CB1Rs are expressed on striatal PV-FSIs and mediate a form of inhibitory LTD that disinhibits MSNs, a mechanism that is associated with striatal regulation of behavioral flexibility (DePoy et al., 2013; Brian N. Mathur et al., 2013). CB1Rs are also expressed on cortical inputs that target MSNs and MSNs themselves (Gerdeman and Lovinger, 2001; Gerdeman et al., 2002; Wu et al., 2015; Lovinger and Mathur, 2012; Lovinger et al., 2022), but it has not yet been established whether cortical projections targeting PV-FSIs also contain CB1Rs. CB1R signaling at cortical-striatal FSI synapses would be expected to reduce inhibitory tone and increase DMS MSN activation, a similar result to CB1R signaling at FSI-MSN synapses. Direct manipulation of DLS PV-FSIs shows that their activity is critical to supporting habitual responding (O’Hare et al., 2017; Patton et al., 2020) but much less is known about DMS PV-FSIs and their contribution to habitual or goal-directed responding. Thus, these two hypotheses must be tested further to discover the mechanism of DMS CB1R regulation of Pavlovian devaluation sensitivity.

Overall, the current studies show that males are sensitive to Pavlovian outcome devaluation, a result that may be explained by reduced inhibitory synaptic transmission in the DMS. We find that the devaluation sensitivity of male rats requires DMS CB1R, but more work is needed to identify the cell-type specific population of CB1Rs that support flexible responding. Additionally, it is possible that DMS CB1Rs would be necessary for the devaluation sensitivity of females in cases where they respond flexibly at baseline. Thus, future studies should manipulate DMS endocannabinoids under conditions in which males and females respond similarly to determine if CB1Rs play a sex-specific role in mediating behavioral flexibility.

## Notes

### Competing Interest Statement

The authors have declared no competing interest.

### Summary of Updates

Materials and methods updated to include the description of slice electrophysiology methods. Results updated to clarify male specific findings in outcome devaluation. Results extended to include slice electrophysiology findings (Figure 4 and 5). Figure 2 is revised to only show overall outcome devaluation findings. Figure 4 from previous version is moved to supplemental data and we also include additional supplemental data (Fig. 2-1, 2-2,3-1 and 3-2). Discussion is expanded upon to synthesize behavioral experiments and slice electrophysiology findings. Author list added to include additional collaborator.

