## Supplemental Figures for "Dorsomedial Striatum CB1R signaling is required for Pavlovian outcome devaluation in male Long Evans rats and reduces inhibitory synaptic transmission in both sexes"

### Slide 1
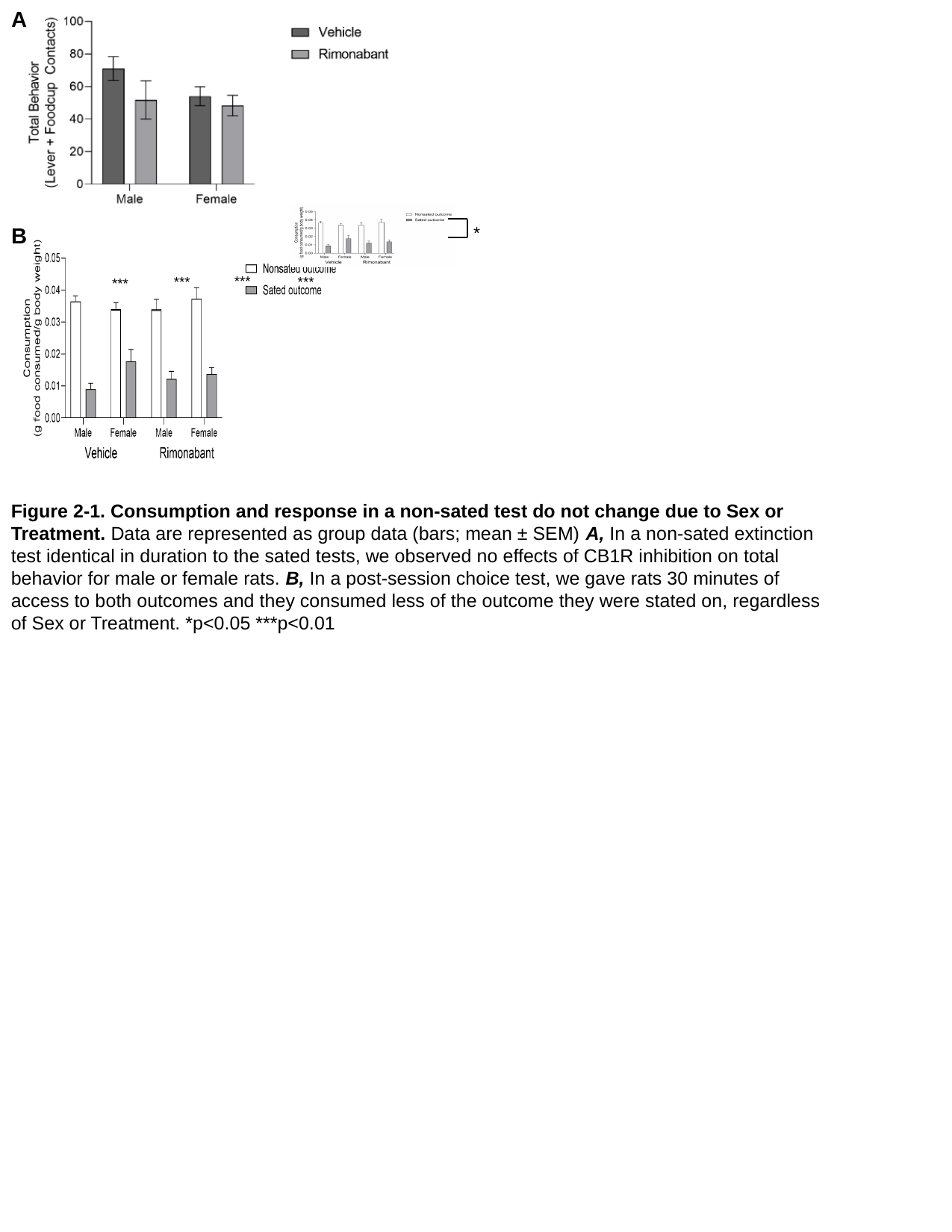

A
***
***
***
***
*
B
Figure 2-1. Consumption and response in a non-sated test do not change due to Sex or Treatment. Data are represented as group data (bars; mean ± SEM) A, In a non-sated extinction test identical in duration to the sated tests, we observed no effects of CB1R inhibition on total behavior for male or female rats. B, In a post-session choice test, we gave rats 30 minutes of access to both outcomes and they consumed less of the outcome they were stated on, regardless of Sex or Treatment. *p<0.05 ***p<0.01

### Slide 2
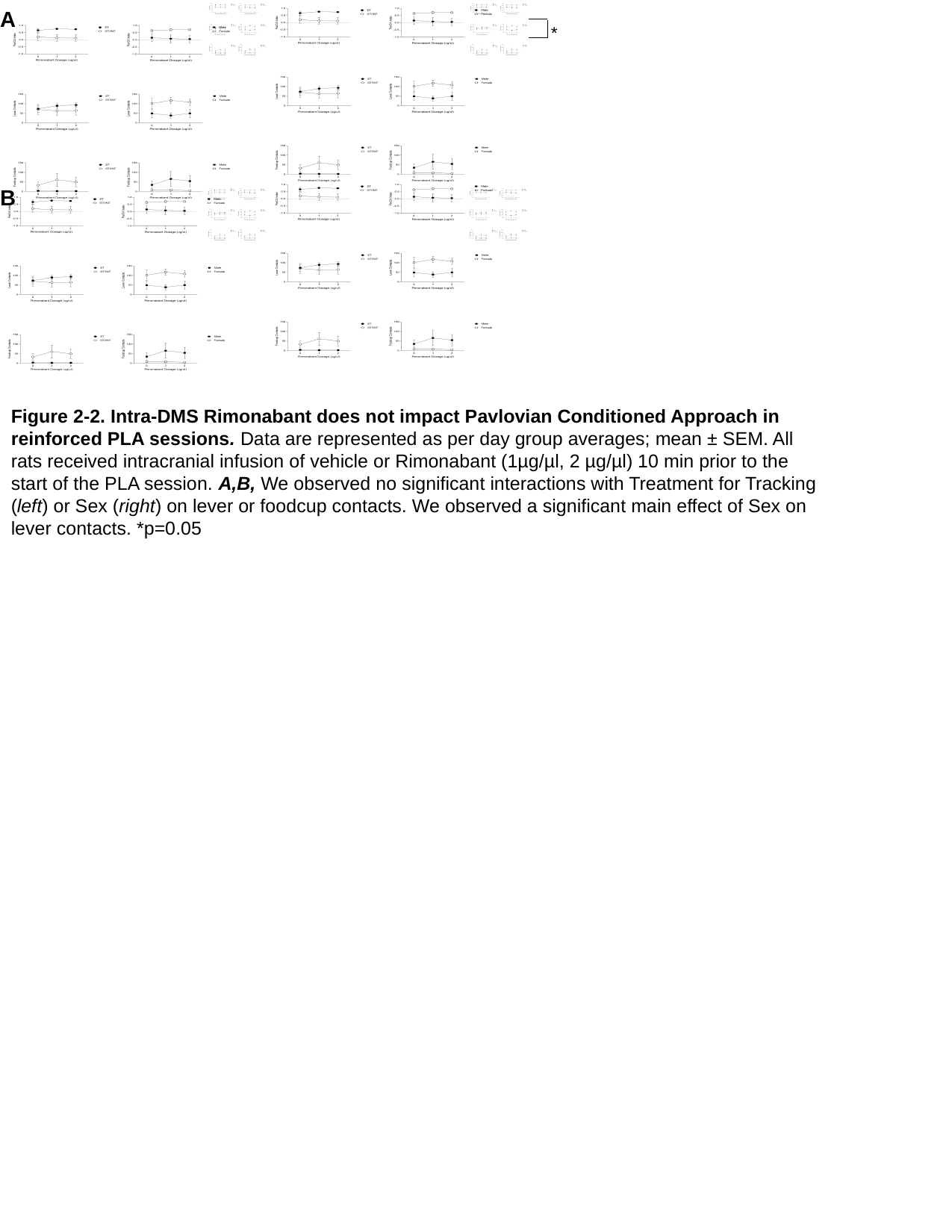

A
*
B
Figure 2-2. Intra-DMS Rimonabant does not impact Pavlovian Conditioned Approach in reinforced PLA sessions. Data are represented as per day group averages; mean ± SEM. All rats received intracranial infusion of vehicle or Rimonabant (1µg/µl, 2 µg/µl) 10 min prior to the start of the PLA session. A,B, We observed no significant interactions with Treatment for Tracking (left) or Sex (right) on lever or foodcup contacts. We observed a significant main effect of Sex on lever contacts. *p=0.05

### Slide 3
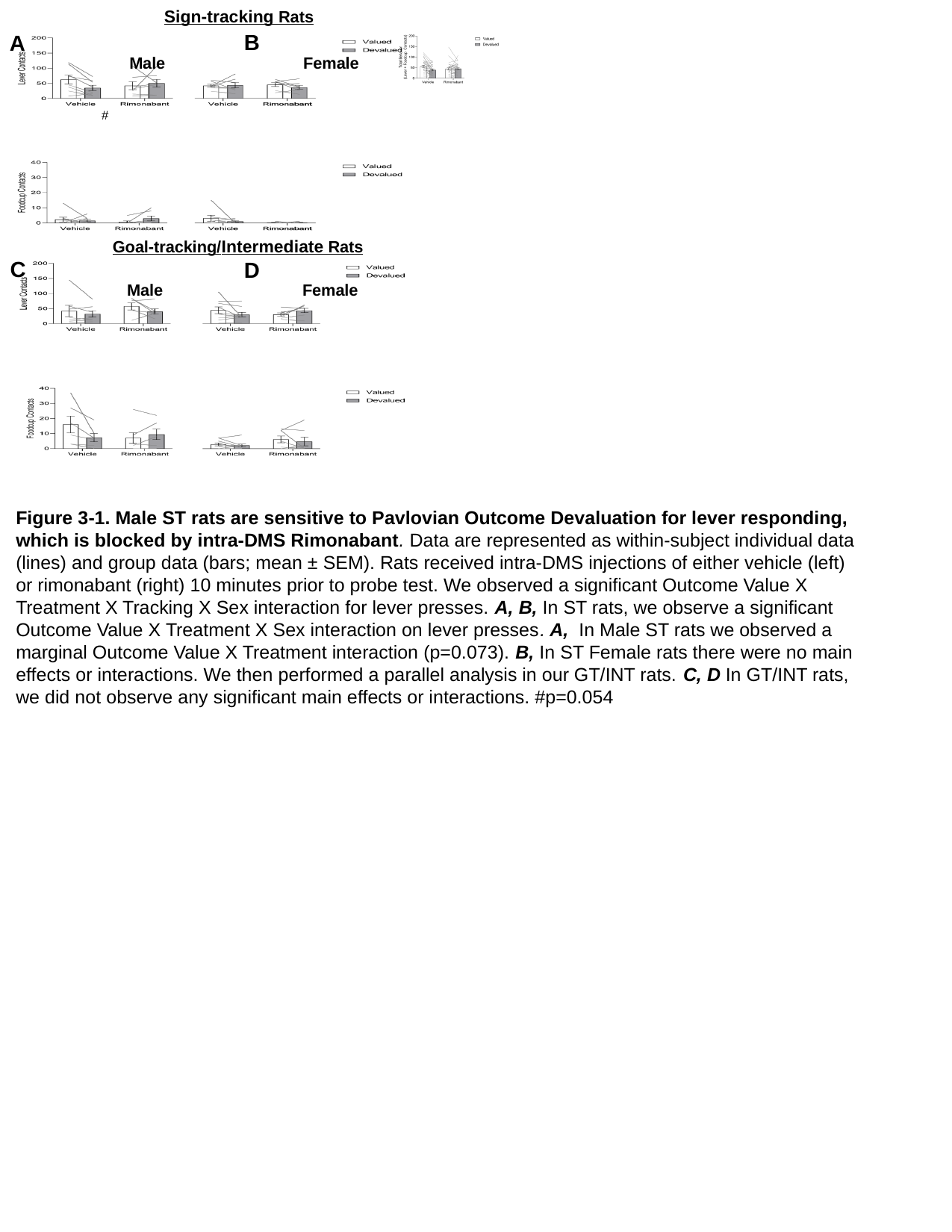

Sign-tracking Rats
B
A
#
Female
Male
Goal-tracking/Intermediate Rats
C
D
Male
Female
Figure 3-1. Male ST rats are sensitive to Pavlovian Outcome Devaluation for lever responding, which is blocked by intra-DMS Rimonabant. Data are represented as within-subject individual data (lines) and group data (bars; mean ± SEM). Rats received intra-DMS injections of either vehicle (left) or rimonabant (right) 10 minutes prior to probe test. We observed a significant Outcome Value X Treatment X Tracking X Sex interaction for lever presses. A, B, In ST rats, we observe a significant Outcome Value X Treatment X Sex interaction on lever presses. A, In Male ST rats we observed a marginal Outcome Value X Treatment interaction (p=0.073). B, In ST Female rats there were no main effects or interactions. We then performed a parallel analysis in our GT/INT rats. C, D In GT/INT rats, we did not observe any significant main effects or interactions. #p=0.054

### Slide 4
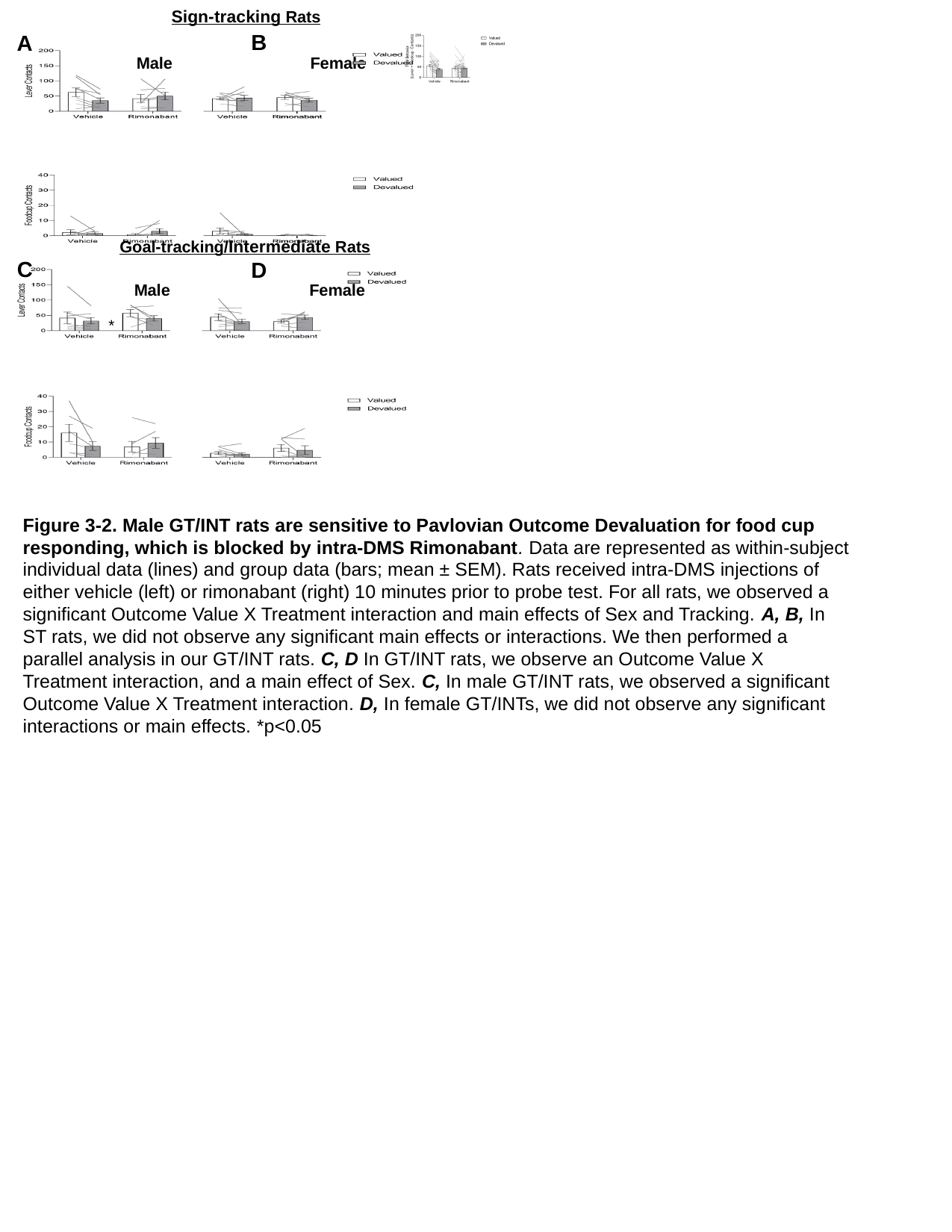

Sign-tracking Rats
B
A
Female
Male
Goal-tracking/Intermediate Rats
C
D
Male
Female
*
Figure 3-2. Male GT/INT rats are sensitive to Pavlovian Outcome Devaluation for food cup responding, which is blocked by intra-DMS Rimonabant. Data are represented as within-subject individual data (lines) and group data (bars; mean ± SEM). Rats received intra-DMS injections of either vehicle (left) or rimonabant (right) 10 minutes prior to probe test. For all rats, we observed a significant Outcome Value X Treatment interaction and main effects of Sex and Tracking. A, B, In ST rats, we did not observe any significant main effects or interactions. We then performed a parallel analysis in our GT/INT rats. C, D In GT/INT rats, we observe an Outcome Value X Treatment interaction, and a main effect of Sex. C, In male GT/INT rats, we observed a significant Outcome Value X Treatment interaction. D, In female GT/INTs, we did not observe any significant interactions or main effects. *p<0.05
